# Longitudinal to transverse metachronal wave transitions in an in-vitro model of ciliated bronchial epithelium

**DOI:** 10.1101/2021.10.28.466261

**Authors:** Olivier Mesdjian, Chenglei Wang, Simon Gsell, Umberto D’Ortona, Julien Favier, Annie Viallat, Etienne Loiseau

**Affiliations:** Aix Marseille Univ, CNRS, CINAM, Turing Centre for Living systems, Marseille, France; Aix Marseille Univ, CNRS, Centrale Marseille, M2P2, France

## Abstract

Myriads of cilia beat on ciliated epithelia, which are ubiquitous in life. When ciliary beats are synchronized, metachronal waves emerge, whose direction of propagation depends on the living system in an unexplained way. We show on a reconstructed human bronchial epithelium in-vitro that the direction of propagation is determined by the ability of mucus to be transported at the epithelial surface. Numerical simulations show that longitudinal waves maximise the transport of mucus while transverse waves, observed when the mucus is rigid and still, minimize the energy dissipated by the cilia.

---

From swimming microorganisms to major organs, multiciliated cells are ubiquitous in living systems. Active cilia support multiple biological functions such as the motility and feeding of marine microorganisms, circulation of the cerebrospinal fluid in the central nervous system, mucociliary clearance of pathogens and pollutants from airways[1, 2]. The generation of fluid flows at the scale of a microorganism or at the tissue level requires a dense array of cilia that coordinate both their beat directions [3, 4] and their phase [5]. Such coordination results from hydrodynamic coupling between cilia [6]. When neighboring cilia maintain a constant phase difference, metachronal waves emerge [7].

A variety of physical models have been developed to address the conditions of emergence of waves and their possible advantage for biological systems [8–13]. Using an analytical approach, Guirao and Joanny [14] showed that metachronal coordination arises from a selforganized internal phenomenon and hydrodynamic coupling that improves the steadiness of the resulting flow. More sophisticated models based on numerical simulations solve the fluid-structure interaction of flexible cilia beating in fluid [15, 16]. By varying the geometrical arrangement of the cilia, they show that metachrony results in an increased velocity transport and efficiency. Elgeti et al. [15] observe the emergence of waves propagating in various direction: longitudinal, transverse and oblique. Yet, the role of the direction of propagation is not clear.

In marine organisms, ciliated epithelia exhibit metachronal waves propagating in different directions [17, 18], but the majority presents transverse waves. On the contrary, observations made ex-vivo on animal bronchial epithelia report the presence of longitudinal waves [19–21]. These experimental results suggest that the direction of propagation may play a role to optimise a given biological function.

The relationship between the type of metachronism, the biophysical parameter the living system optimizes and the physiological function are still unclear. An experimental system where the emergence of metachrony can be tuned would be a major asset to decipher such relationship.

In this study, we managed to control the type of metachronal waves that emerge at the surface of an invitro reconstituted bronchial epithelium, by tuning the boundary conditions on top of cilia. We show that longitudinal waves propagate when mucus is transported at the epithelial surface, while the presence of a stuck layer of mucus above cilia results in a transition towards transverse metachronal waves. These results are in good agreement with a numerical model built on the lattice Boltzmann Method (LBM) coupled with immersed boundaries to solve the fluid-structure interactions of flexible beating cilia. We show that longitudinal waves are better suited to transport fluid while when a no slip condition is imposed on top of cilia, transverse waves minimise the power dissipated by cilia unable to propel mucus.

## Experimental system

The bronchial epithelium is reconstituted in-vitro at the air liquid interface and is composed of basal cells, goblet cells that produce mucus and ciliated cells. The latter cover 80% of the epithelial surface. The apical surface of a ciliated cell (about 30*µm*^2^) bears up to 200-300 beating cilia [22] of 7 µm height and gathered into 10-20 bundles (see Movie 1). Cilia beat in an aqueous periciliary layer of low viscosity (close to that of water) [23] and propel a layer of mucus at their tips along the epithelial surface (Fig. 1(a)).

**FIG. 1.**
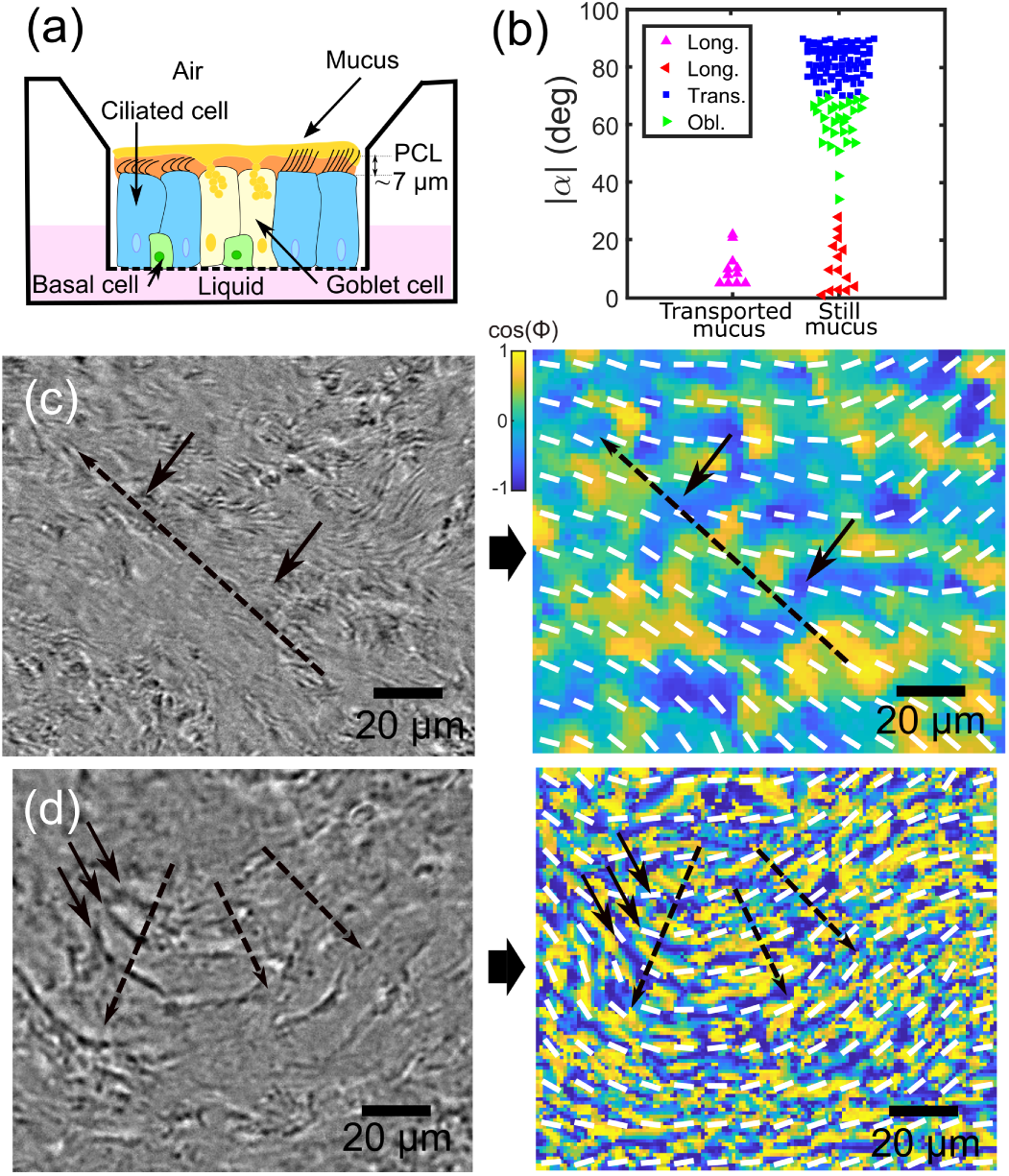
(a) Schematic of an air-liquid culture of a reconstituted human bronchial epithelium. The epithelium is constituted of ciliated cells, goblet cells and basal cells. Cilia beat in the aqueous periciliary layer and their tips poke in the above mucus layer during the forward stroke. (b) Angle | *α*| between the direction of propagation of the metachronal wave and the ciliary beat direction in transported and still mucus condition. (c) Longitudinal wave in the transported mucus condition. (d) Transverse waves in the still mucus condition. On the left, brightfield image of the propagating wave. The direction of propagation is indicated by the dotted arrow, and the wave fronts are indicated by two solid arrows. On the right, corresponding map of *cos*(Φ), with Φ the local phase of beat pattern, obtained by FFT analysis (see Supplemental Material, Fig. 1 for details). The local beat direction is displayed in white.

## Materials and methods

Cultures of human bronchial epithelium reconstituted from primary cells in 6*mm* diameter transwells were purchased from Epithelix (MucilAir). The cultures were maintained in an incubator at 37°*C* and 5% *CO*_2_. Epithelix MucilAir culture medium (700 µl) was replaced every 2 days. The mucus on the apical epithelial surface was either kept moist by adding 3*µL* of culture medium to the surface once a week to compensate for evaporation processes, which was sufficient to maintain a rotating transport of mucus on the epithelial surface (Movie 2, [3, 4]), or allowed to dry spontaneously to gradually form a ‘crust’ of mucus. In the latter case, mucus transport speed gradually decreased until only short-range vibrations were observed (Movie 3), but the beating of the cilia under the mucus layer could still be observed. In total, we worked on 4 wells with moistened mucus from 3 donors and 5 wells with dried mucus from 3 donors.

Cultures were observed in brightfield on a Nikon Eclipse Ti inverted microscope with a 20x objective and 1.5x magnification lens, at 37°*C* under humidified airflow with 5% *CO*_2_ with a Photron camera (Fastcam, mini UX50) at 250 fps or a Luminera camera (Infinity 3 Luminera USB) at 40 fps. We performed a gentle apical whasing before imaging by adding and removing 200 µl of culture medium.

## Emergence of metachronal waves at the epithelial surface

At the scale of a ciliated cell, we observe bundles of cilia that beat asymmetrically at a frequency ranging between 7 and 12*Hz* (see Supplemental Material, Fig. 2). On a single cell, we observe that the bundles have the same frequency and the same beating direction. Bundles aligned along the beating direction beat synchronously over a typical length scale of a ciliated cell (∼5 − 6*µm*). In the direction perpendicular to the beating direction, phase shifts between the beats of the different bundles are observed, which become significant at the scale of half a cell (see Movie 1). Indeed, the distance between neighboring bundles on the apical surface of the same cell being less than the length of a cilia, steric constraints impose synchronization along the stroke direction.

At the tissue scale, we observe the propagation of localized metachronal waves on the epithelial surface. The typical distance of propagation is of the order of 40*µm*, i.e. over approximately 8 cells and the width of the wave-front ranges from 20 to 40*µm* (see Fig. 1(c) and 1(d)). The metachronal waves are observed regardless of the collective direction of the ciliary beats, whether linear or circular (See Movie 6).

We observe three different types of metachronal waves characterized by the angle *α* between the direction of propagation and the ciliary beating direction (see Fig. 1(b)). The first type gathers longitudinal waves (|*α*| < 15°), which propagate in the same direction than the forward stroke (symplectic waves). Interestingly, this type of metachrony is the only one that emerges on cultures where the mucus is continuously transported (Fig. 1(b) and (c), Movies 4-5). On rare occasions, symplectic waves emerge on cultures with dried and still mucus (Fig. 1(b)). The two other types of metachronal waves are observed exclusively on cultures with dried still mucus: (i) transverse waves which propagate perpendicularly to the direction of the ciliary beats (Fig. 1(c) and (b), Movies 6-7-8), and (ii), oblique waves that propagate along a direction that makes an angle |*α*| between 30° and 70° with regards to the beating direction (Fig. 1(b), Movie 9).

We measured the wavelength and the distance of wave propagation on each culture. The distributions are displayed in figure 2. The average wavelength of longitudinal waves is 40*µm*, i.e. 8 cells long. While transverse and oblique waves exhibit the same wavelength, much shorter, *λ*_*w*_ = 13*µm*, about only 2 ciliated cells long (Fig. 2(a)). The distance of propagation is of the same order of magnitude for the three types of wave, in the range of 15-60 µm (Fig. 2(b)). It is worth noting that for a wave to be able to propagate, all cilia must beat at the same frequency. We established frequency maps and identified domains with equal beat frequencies (see Fig. 2(c)), whose size limits the distance of wave propagation. We found that the size of these domains is in the range of 15-60 µm (see Fig. 2(d)), that corresponds well to the propagation distance of the metachronal waves. Note that the same size distribution has been established in the dried mucus condition (see Supplemental Material, Fig. 3). The ratio between the distance of propagation and *λ*_*w*_ corresponds to the number of wave front that can be observed in the system. It is of the order of 1 for longitudinal waves and can reach up to 6 fronts for oblique and transverse waves (see Supplemental Material, Fig. 2).

**FIG. 2.**
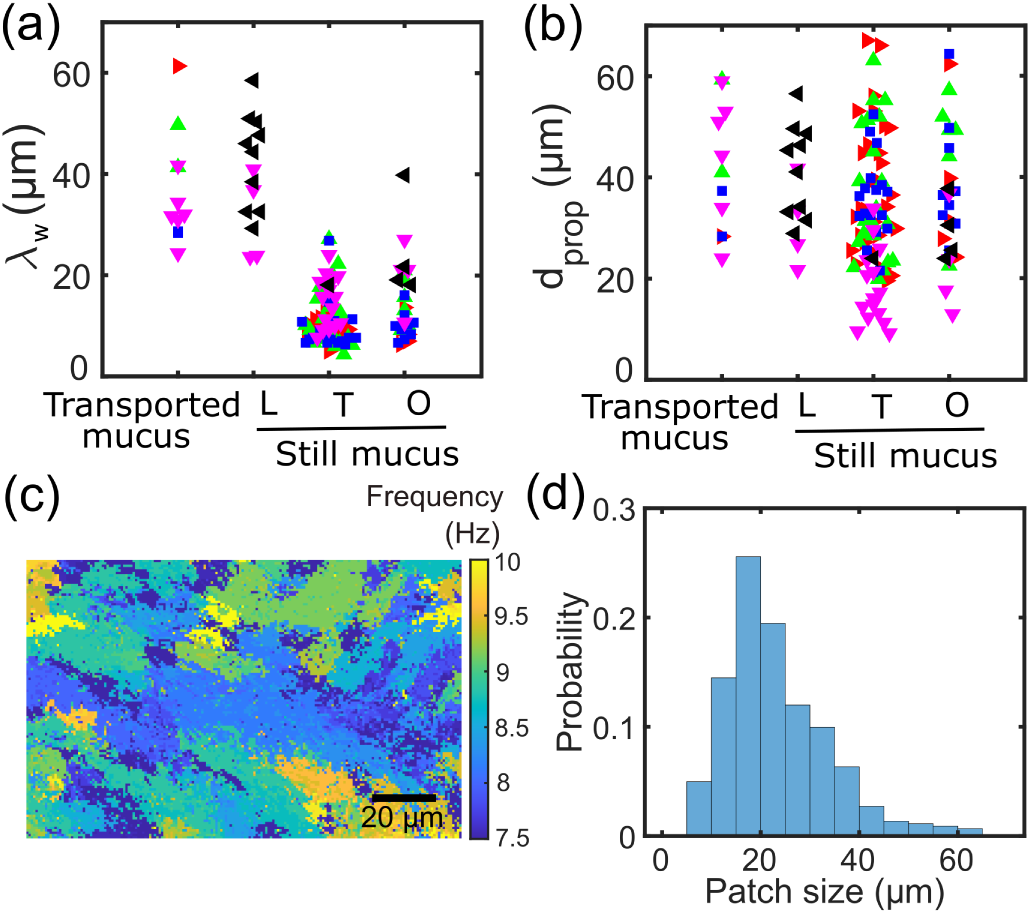
(a) Distributions of wavelengths λ_*w*_ and (b) distance of propagation for longitudinal (L), transverse (T) and oblique (O) waves in transported mucus and still mucus conditions. Each color represents a different culture, N=4 cultures in the transported mucus condition, and N=5 cultures in the still mucus condition. (c) Map of ciliary beat frequencies computed with FFT analysis (see Supplemental Material Fig. 1 for details) in the transported mucus condition. (d) Size distribution of iso-frequency patches. The total area selected for the measurements is 420*µm ×* 340*µm*.

Experimentally, on reconstituted epithelia, we observed longitudinal waves associated with *λ*_*w*_ = 40*µm*. When the mucus is dried and not transported anymore, transverse and oblique waves emerge associated with a shorter wavelength (*λ*_*w*_ = 13*µm*). To understand the mechanisms at the origin of the emergence of the different observed metachronal waves, we developed a numerical hydrodynamic model of flexible cilia.

## The model

We modelled a cilium as a slender elastic body of diameter D and length L, with *D* ≪ *L*. The internal actuation of each cilium is set by its basal angle *β*, which is forced to vary in a wave-like form in space and time, with a pulsation *ω* and a wavelength *λ*:

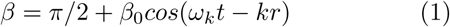

with *ω*_*k*_ = *π/T*_*k*_ where *T*_*k*_ = *T*_*p*_ during the power stroke and *T*_*k*_ = *T*_*r*_ during the recovery stroke, *k* = 2*π/λ* and *r* = *n*Δ with *n* = 1, 2, 3, … and Δ is the distance between two cilia. The cilium dynamics is governed by the equation of elasticity described in [24], with a time dependent bending rigidity. We obtained a two-dimensional beating pattern for one cilium in agreement with those described in the literature [19] (see Fig. 3(a)). The forward stroke is in the increasing x direction. The dynamics of the three-dimensional incompressible flow described by the continuity and momentum equations, is solved using the lattice Boltzmann method, following [16]. The fluid is considered as Newtonian with a Reynolds number fixed to 0.5. To account for the unsteady flexible boundaries, we use the immersed boundary method (IBM), following the numerical framework proposed in [24]. The details of the method for the numerical simulation is in the SI. Cilia are placed along a line in the numerical domain and distant of Δ = 0.5*L* or Δ = 1*L* depending on the boundary conditions. The transverse waves are obtained by setting the cilia along the y axis (see Fig. 3(b)), whereas the longitudinal waves are obtained by setting the cilia along the x axis (see Fig. 3(c)). The height of the domain *h* is fixed to 1.2*L*. On top of cilia, we modelled the flowing mucus condition or the stuck mucus condition by either a slip or no slip condition for the fluid. The wavelength *λ* is fixed by the number of cilia present in the domain, and the periodic condition is imposed on both the x and y direction. We computed several quantities related to the cilia and the flow.

**FIG. 3.**
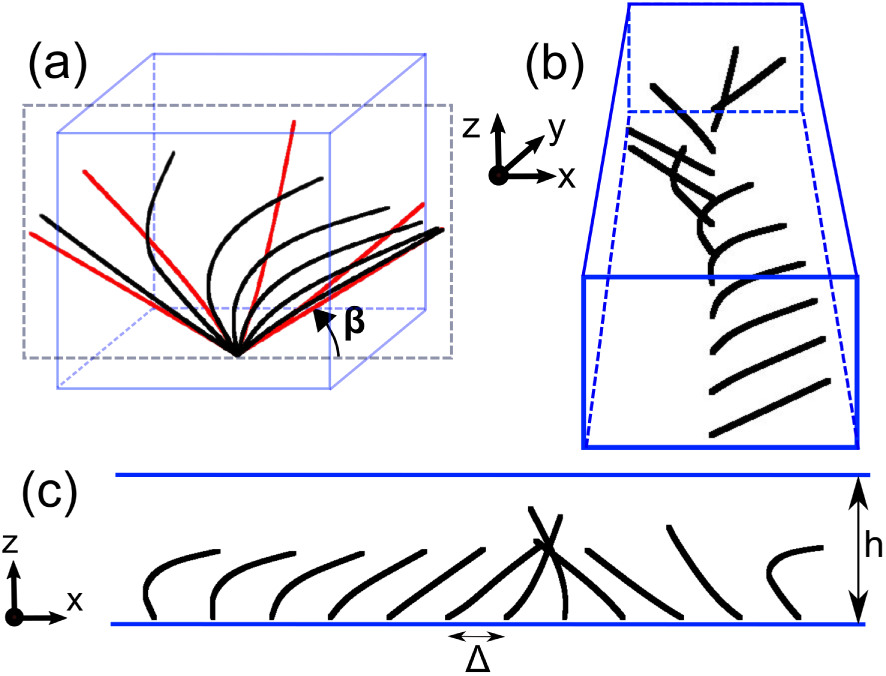
Geometrical configurations used in the model for longitudinal and transverse waves. (a) Beat pattern of a cilium of length L. In red succesive positions during the forward stroke and in black during the recovery. *β* is the basal angle (see main text). The cilium motion stays in the *y* = 0 plane (black dotted rectangle) with the power stroke oriented towards the increasing x direction. (b) Cilia aligned in the y direction: example of a transverse configuration with *λ* = 12*L* and Δ = 1*L*. (c) Cilia aligned in the x direction: example of a longitudinal configuration with *λ* = 12*L* and Δ = 0.5*L*. The height of the domain is fixed to *h* = 1.2*L*.

The non-dimensional time-dependent force and power that one cilium exerts on the fluid are defined respectively as 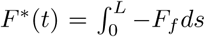 and 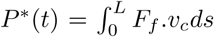, with *F*_*f*_ the force that the fluid exerts on the cilium, and *V*_*c*_ the velocity of the cilium. We converted *F*^*^ and *P*^*^ into their respective dimensional quantities *F* = *F*^*^ ×(*EI/L*^2^) and *P* = *P*^*^ × (*V*_*c*_*EI/L*^2^), with *EI* is the bending rigidity of one cilium, *L* the length of one cilium, and *V*_*c*_ = 2*L/T*_*b*_ the characteristic velocity of one cilium, with *T*_*b*_ = *T*_*r*_+*T*_*p*_ the beating period. We used the experimental values found in the literature *EI* = 6.10^−22^*N*.*m*^2^ [25], *L* = 7*µm* and *T* = 1*/*10*s* [19].

As shown in the Supplemental Material Fig. 4, *P*(*t*) exhibits periodically two regions: a first long duration with a small peak corresponding to the cilium recovery stroke beating, and a second shorter duration with a higher peak corresponding to the cilium power stroke beating. We note 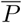 (resp. 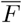) the average of *P*(*t*) (resp. *F*(*t*)) on one beating period at steady state.

We also computed the time dependent flux generated by one cilium in the x direction (resp. y direction) crossing the yz plane (resp. the xz plane) 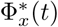 (resp. 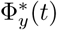), whose relations with the dimensional flux are respectively 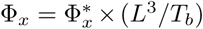 and 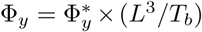. These fluxes typically reach a steady state after the first half period of cilia beating (see Supplemental Material, Fig. 4). We deduced the total mean flux generated by one cilium as 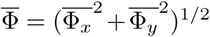, with 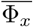 (resp. 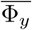) the time average of Φ_*x*_ (resp. Φ_*y*_) over one period at steady state.

Finally we defined the mean fluid velocities as 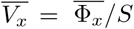 and 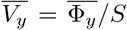 where *S* is the lateral section of the domain corresponding to one cilium *S* = *h* × Δ = (1.2*L*) × Δ, with Δ = 0.5*L* or Δ = 1*L*.

## Slip condition : Efficiency of longitudinal metachronal waves

Experimental observations show that only longitunal waves propagate. In the model, we set Δ = 1*L*, since the length of one cell is approximately the same as one cilium and that at the scale of one multiciliated cells cilia beat synchronously. We chose to compute the mucus flux based on the reasonable hypothesis that bundles of cilia synchronize themselves in order to maximize mucus transport. Figure 4(a) shows the variation of the flux with the wavelength for both longitudinal and transverse waves. The flux created by the longitudinal waves becomes higher than the flux of the transverse waves for wavelengths larger than ∼50*µm*, in the range of the observed longitudinal wavelength (Fig. 2(a)). In principle, we should expect that waves of higher wavelength should be observed since the flux increases with the wavelength. However, the biological system prevents the emergence of such waves due to the limited size (50*µm*) of multiciliated cell domains with equal frequency.

**FIG. 4.**
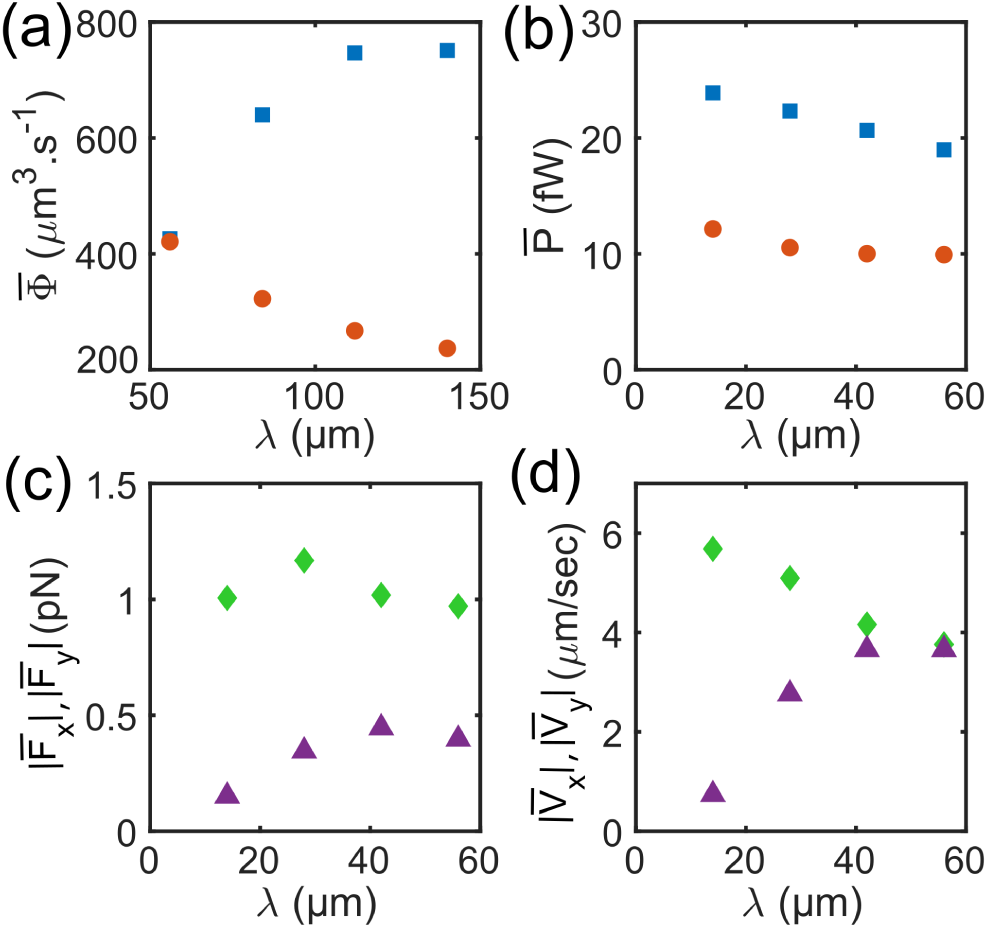
Averaged quantities numerically computed and plotted as a function of *λ*. (a) Total mean flux for longitudinal (blue square) and transverse (red circle) waves in the slip condition with Δ = 1*L*. (b) Mechanical power dissipated by one cilium for longitudinal (blue square) and transverse (red circle) waves in the no slip condition with Δ = 0.5*L*. In the case of transverse waves, in no slip condition and Δ = 0.5*L*, (c) force exerted by a cilium along x (green diamond) and along y (purple triangle), and (D) local fluid velocity at a cilium position, along x (green diamond) and along y (purple triangle).

## No slip condition : power dissipation and transverse flow

Experimental observations show the emergence of transverse waves in absence of mucus transport. In this condition, as phase shift are observed at the scale of half a cell. We set Δ = 0.5L in the model. As the flux is no longer a relevant quantity, we hypothetize that bundles synchronize themselves to minimize the dissipated mechanical power. Figure 4(b) shows the variation of the dissipated power with the wavelength. At all wavelengths 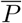 of transverse waves is about two times lower than longitudinal waves. This explains the observed emergence of transverse waves in absence of mucus transport. However, there is no strong variation of 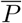 with *λ* for transverse waves which could explain why short wavelengths are experimentally observed. In order to have a deeper insight in the sytem, we computed the hydrodynamic force and the fluid velocity in the PCL exerted by a cilium. The x and y components of these quantities are shown in Fig. 4(c) and 4(d), respectively. Strikingly, the y-component of 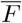 increases by a factor 3 and by a factor of 8 for 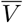 between *λ* = 14*µm* and *λ* = 42*µm*. The x-component of 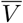 decreases from 6 to 4*µm/sec* between *λ* = 14*µm* and *λ* = 42*µm* so that at *λ* = 42*µm*, 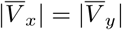. At all *λ*, 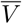 is of the same order of magnitude but its direction is progressively shifted from a longitudinal direction for *λ* = 14*µm* towards 45° for *λ* = 42*µm*. The flow can be moved even more in the transverse direction if the cilia are set closer than Δ = 0.5*L*, as showed in the Supplemental Material, Fig. 5. We consider that such a flow is susceptible to affect the direction of neighboring ciliary beat. Indeed, cilia are mechanosensitive systems which can modify their beat characteristics in response to an external hydrodynamic cue [3, 26, 27]. Based on these considerations, the numerical model suggests that the system favours the emergence of transverse waves with short wavelength to prevent a possible destabilisation of ciliary beat directions. This corresponds to the previously described experimental observations.

## Concluding remarks

We experimentally showed that the type of metachronism could be modified on the same system and depends on the external conditions. The combination of experimental and numerical approaches revealed the system optimises the flux to transport the mucus, while it lowers the energy consumption and prevent any hydrodynamic destabilisation of the cilia in absence of transport. This approach on a biological system of great importance clearly shows that each type of metachronism could be associated with the optimisation of a physical quantity, linked with a physiological function or regulation of the biological system. Our study may shed new light on metachronisms in other living sys-tems such as marine organisms.

## ACKNOWLEDGMENTS

We thank Dr. K. Khelloufi for providing a movie. The project leading to this publication has received funding from the “Investissements d’Avenir” French Government program managed by the French National Research Agency (ANR-16-CONV-0001) and from Excellence Initiative of Aix-Marseille University - A*MIDEX. The Centre de Calcul Intensif d’Aix-Marseille University is acknowledged for granting access to its high performance computing resources.

